# Kinetic models of metabolism that consider alternative steady-state solutions of intracellular fluxes and concentrations

**DOI:** 10.1101/437822

**Authors:** Tuure Hameri, Georgios Fengos, Meric Ataman, Ljubisa Miskovic, Vassily Hatzimanikatis

**Affiliations:** Laboratory of Computational Systems Biotechnology (LCSB), Swiss Federal Institute of Technology (EPFL), CH-1015 Lausanne, Switzerland.

## Abstract

Large-scale kinetic models are used for designing, predicting, and understanding the metabolic responses of living cells. Kinetic models are particularly attractive for the biosynthesis of target molecules in cells as they are typically better than other types of models at capturing the complex cellular biochemistry. Using simpler stoichiometric models as scaffolds, kinetic models are built around a steady-state flux profile and a metabolite concentration vector that are typically determined via optimization. However, as the underlying optimization problem is underdetermined, even after incorporating available experimental omics data, one cannot uniquely determine the operational configuration in terms of metabolic fluxes and metabolite concentrations. As a result, some reactions can operate in either the forward or reverse direction while still agreeing with the observed physiology. Here, we analyze how the underlying uncertainty in intracellular fluxes and concentrations affects predictions of constructed kinetic models and their design in metabolic engineering and systems biology studies. To this end, we integrated the omics data of optimally grown *Escherichia coli* into a stoichiometric model and constructed populations of non-linear large-scale kinetic models of alternative steady-state solutions consistent with the physiology of the *E. coli* aerobic metabolism. We performed metabolic control analysis (MCA) on these models, highlighting that MCA-based metabolic engineering decisions are strongly affected by the selected steady state and appear to be more sensitive to concentration values rather than flux values. To incorporate this into future studies, we propose a workflow for moving towards more reliable and robust predictions that are consistent with all alternative steady-state solutions. This workflow can be applied to all kinetic models to improve the consistency and accuracy of their predictions. Additionally, we show that, irrespective of the alternative steady-state solution, increased activity of phosphofructokinase and decreased ATP maintenance requirements would improve cellular growth of optimally grown *E. coli*.

## 1. Introduction

Over the last decades, advances in genome editing technologies have allowed the redirection of carbon flow within the organism towards specialty products of interest and desired physiologies [1]. Identifying candidate enzymes is fundamental for genetic modifications that have seen applications in metabolic engineering, basic and applied biology, biotechnology and medical sciences [2–4]. Increasingly available high-throughput sequencing data has enabled the construction of stoichiometric genome-scale metabolic models (GEMs) that describe mathematically the balanced metabolic fluxes within an organism [5]. Metabolic models such as these GEMs have been extensively used to characterize overall network behavior of organisms, which can provide guidance about the genes that can be modified to improve a desired product biosynthesis. Improved guidance for metabolic engineering and basic biology will be achieved with kinetic models of the reactions/networks in GEMs.

The construction of a kinetic model of metabolism requires knowledge of steady states and/or dynamics of metabolic fluxes and metabolite concentrations that can be used to estimate the unknown kinetic parameters that describe these data. However, there are many sources of uncertainty in metabolic fluxes and metabolite concentrations that hamper the accurate estimation of kinetic parameters. Advances in C13 isotopomer techniques facilitated the measurement of fluxes across cellular reactions and promoted the development of metabolic flux analysis (MFA) [6]. One main uncertainty in fluxes is the flux directionality as reactions can be thermodynamically bidirectional [7]. Metabolomics and thermodynamics can be used as it is done in thermodynamic-based flux analysis (TFA) [7–9] to constrain the direction of some of these fluxes. But even when information about the directionality of all the reactions and fluxomics from labeling experiments are used, there is still a great uncertainty on exact estimation of fluxes as the degrees of freedom remain high, especially as we increase the size of the networks. The addition of constraints based on measured gene expression data [10, 11] and enzymatic data [12] can reduce the degrees of freedom. However, the system remains underdetermined, resulting in multiple alternative steady-state flux distributions corresponding to the physiology under study. Different steady-state solutions could directly affect the predictions of kinetic models, leading towards very distinct conclusions and guidance for metabolic engineering.

Several promising methods exist for constructing kinetic models around representative steady states of metabolic fluxes and metabolite concentrations [13, 14]. The Optimization and Risk Analysis of Complex Living Entities (ORACLE) workflow [15–17] and frameworks built around ensemble modeling [18, 19] have made significant strides towards genome-scale kinetic modeling of metabolism. These methods generate populations of non-linear kinetic models around a selected reference steady state (RSS) that is chosen based on its ability to characterize the observed physiology. Methods commonly used for selecting a RSS include using the computed optimal solution to an objective function that defines physiological tasks [20], fitting the data from MFA [6], or performing principal component analysis (PCA) on a sampled solution space [15]. Once a RSS is established, kinetic models are constructed around it, which allows the study and prediction of cellular metabolic response to perturbations [21]. These populations of kinetic models can be studied using statistical procedures to identify target enzymes, sensitively analyze kinetic parameters, and design experiments [22, 23]. There is no unique and evident approach for selecting a RSS for such an underdetermined system. To our knowledge, the impact of alternative RSSs describing a physiology using the kinetic parameters and the outputs of these kinetic models have not been studied.

Hereby, we examine how uncertainty in intracellular flux solutions and metabolite concentrations influences the metabolic control analysis (MCA) of populations of non-linear kinetic models built around alternative steady states. We integrated physiological data from *E. coli* grown aerobically in a batch cultivation [24] into a reduced core model derived from the iJO1366 *E. coli* GEM [25–27] and found that the data were not sufficient to uniquely determine the steady state metabolic flux distribution as several reactions could operate in either the forward or reverse direction. These so-called bi-directional reactions result in the existence of multiple feasible flux directionality profiles (FDPs) that represent the same physiology, because in any FDP, reactions operate only in one direction [8]. We constructed populations of kinetic models for 4 selected FDPs to demonstrate how significantly MCA outputs and metabolic engineering decisions are affected.

## 2. Results and Discussion

The procedure for characterization and analysis of steady-state multiplicities arising from the underdetermined nature of the system is a constitutive part of the ORACLE workflow [15–17, 23, 28–30]. The workflow assists with more reliable and robust MCA-based metabolic engineering decisions that will enable the identification of study-specific target enzymes, independent of the steady state. Various types of biological data are combined into a thermodynamically feasible stoichiometric model of a given physiology (Figure 1). We follow this workflow to discuss our results. At first, we identify the bi-directional reactions and determine feasible flux directionality profiles (FDPs). We discuss how alternative FDPs affect the conclusions of kinetic models. We then consider how the flux values and the metabolite concentration levels within a FDP affect kinetic model predictions. The MCA outputs of the kinetic models are studied to systematically derive metabolic engineering decisions. For further information on the methodologies used, we refer the reader to the methods section of the manuscript.

**Figure 1.**
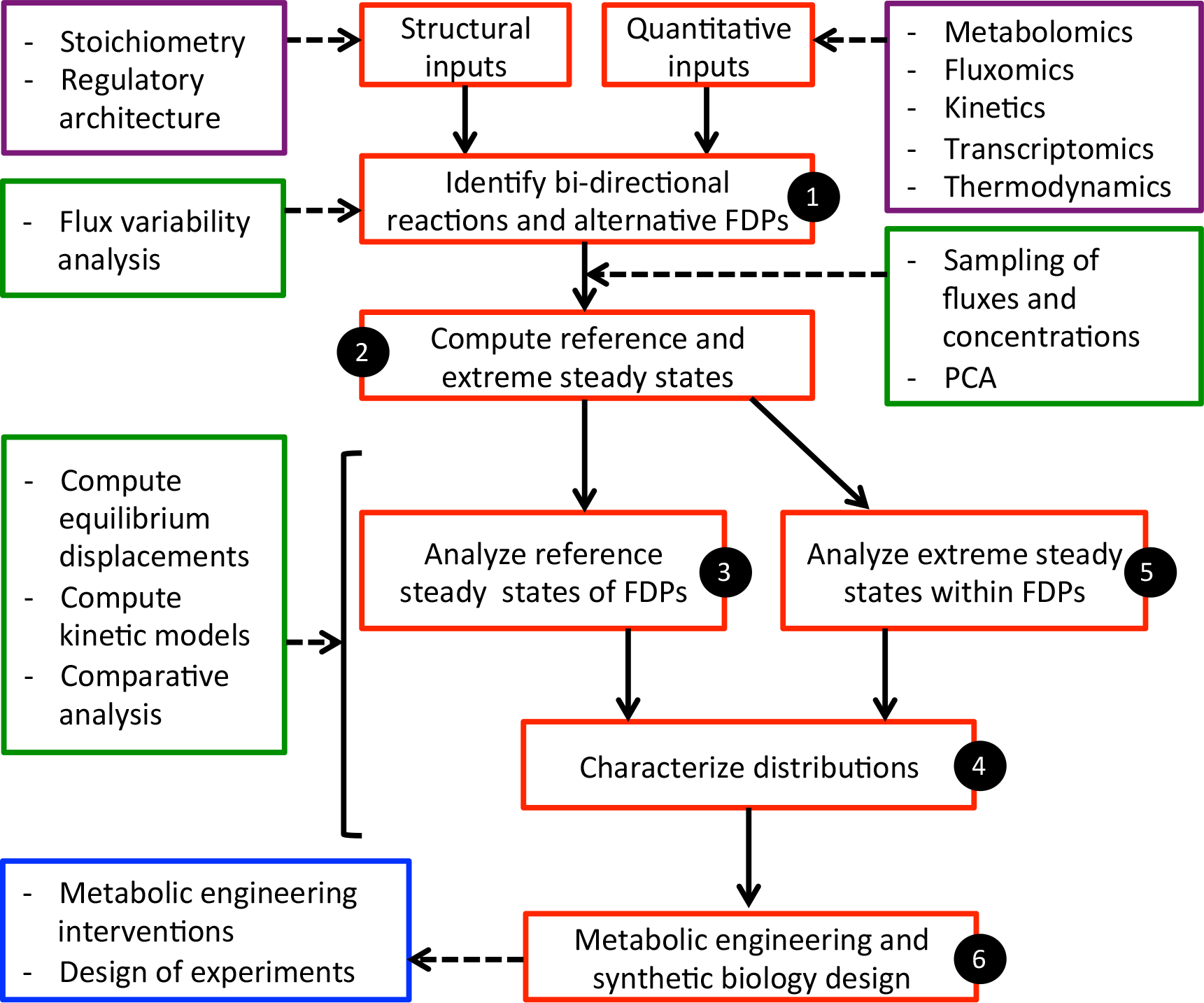
Procedure for characterizing and analyzing multiplicities in metabolic networks. The procedure consists of several computational steps wherein the available data are integrated, the alternative solutions are identified, the populations of non-linear models are built, and the output variables are analyzed to make robust conclusions (for details see main text).

### 2.1. Multiplicity of flux directionality profiles

To derive a reduced *E. coli* metabolic model from the iJO1366 GEM [25], we used the redGEM and lumpGEM algorithms as they provide a systematic and modular way for reducing GEMs, whilst preserving growth and gene essentiality [26, 27]. The obtained core stoichiometric model of the *E. coli* metabolism consisted of 277 reactions and 160 metabolites distributed over the cytosol and the extracellular space (Methods). To constrain the model and derive alternative steady states, we integrated fluxomics and metabolomics data [24] (Supp 1) within the thermodynamic formulation of the stoichiometric model for aerobically grown *E.coli*. Hence, we set the glucose uptake to 7.54 mmol/gDW/h, the growth rate to 0.61 /h and, the excretions of acetate, formate and succinate to 3.5 mmol/gDW/h, 0.5 mmol/gDW/h and 10^−4^ mmol/gDW/h, respectively (Figure 2A). We made assumptions about reaction directionalities based on available literature [24, 31–34] (Methods). Thermodynamic-based variability analysis (TVA) [35] suggested the presence of seven bi-directional reactions in our model: fumarase (FUM), triose-phosphate isomerase (TPI), ribulose-5-phosphate 3-epimerase (RPE), transaldolase (TALA), transketolase 1 (TKT1), transketolase 2 (TKT2), and glucose-6-phosphate isomerase (PGI). All combinations of these seven reactions operating in one or the opposite direction could theoretically lead to up to 128 (2^7^) FDPs. However, due to the stoichiometric and thermodynamic coupling in the network, only 25 out of 128 FDPs were feasible. Some of these reactions such as PGI and FUM are commonly considered as unidirectional. However, Rabinowitz and coworkers reported that these seven identified reactions are bi-directional in *E.coli*, yeast, and immortalized baby mouse kidney cells [36]. This suggests that, for previously uncharacterized physiologies and/or for reactions with no fluxomics data, we should consider all feasible reaction directionalities. This is a way of ensuring that we account for the flexibility of cellular metabolism. For simplicity and clarity of further discussion, we wanted to analyze four FDPs with the most distinct physiologies out of 25 feasible ones. We assumed that changing the directionality of reactions with the largest TVA flux range would result in the most distinct FDPs. PGI and FUM had the largest feasible TVA flux ranges from the seven bi-directional reactions. Hence, to generate these four distinct FDPs, we were changing the directionality of both PGI and FUM in either the forward or backward direction while keeping the directionalities fixed for the 5 remaining bi-directional reactions (Figure 2A). The directionality of these 5 bi-directional reactions (other than PGI and FUM) was determined as follows. We first defined and calculated the flux variability score for each of the 25 FDPs (Methods). A higher flux variability score suggests that the reactions of the FDP are on average more flexible and can operate in relatively wider flux ranges. We then took the directionalities of the remaining five bi-directional reactions from the FDP with the highest score. In this study, we assessed the model predictions and their implications on metabolic engineering decisions around these four FDPs. Nevertheless, different study-dependent criteria for selecting the FDPs could be devised.

**Figure 2.**
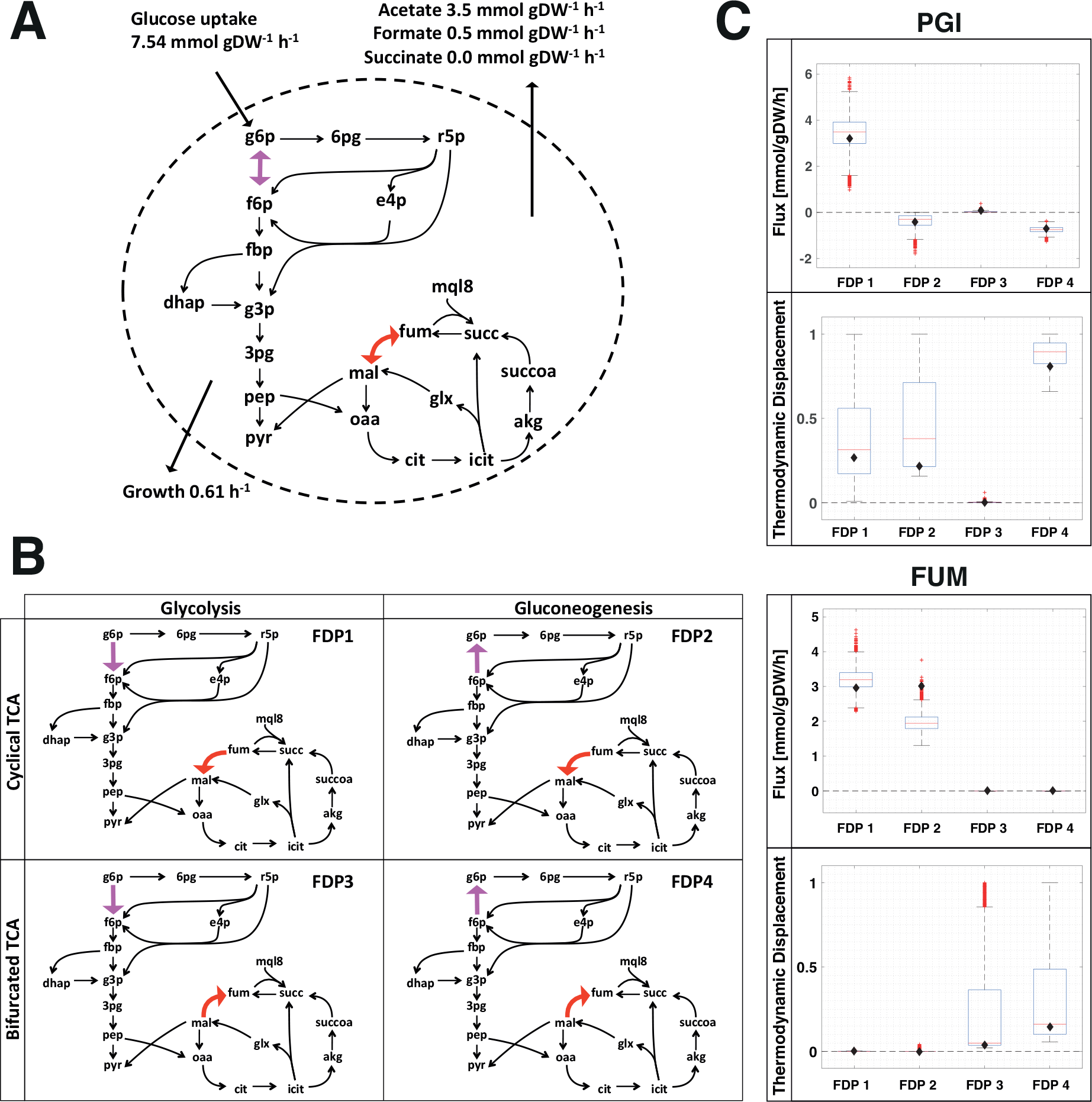
Multiple operational configurations for the same observed physiology of aerobically grown *E. coli*. **(A)** Representation of *E. coli* network. The fluxomics data that were integrated are indicated as uptake, secretion and growth rates. The bidirectional reactions are colored: phosphoglucose isomerase, (PGI, magenta) and fumarase, (FUM, red). **(B)** Representation of the four FDPs for the physiology under study. **(C)** Flux and thermodynamic displacement distributions of PGI and FUM reactions for each of the four generated FDPs. The boxplots show distributions for 5,000 samples. The central red line indicates the median, and the bottom and top edges of the box indicate the 25^th^ and 75^th^ percentiles, respectively. The whiskers correspond to approximately ± 2.7 σ, which is the standard deviation, or 99.3% coverage if the data are normally distributed. Outliers are the points not covered by the range of the whiskers and are plotted individually using the ‘+’ symbol. The black diamond is the RSS value. Full metabolite names are given in supplementary materials (Supp 2).

### 2.2. Comparative analysis of alternative flux directionality profiles

#### 2.2.1. Reference steady states (RSSs) of FDPs

In building kinetic models, we typically must have steady state flux values and metabolite concentrations around which we construct them. We sampled steady states for the flux values and the metabolite concentrations for each FDP and used principal component analysis (PCA) to select their RSSs (Methods and Supp 4). There were considerable differences in the RSS values for the fluxes and thermodynamic displacements of reactions across the network, particularly for Triosephosphate isomerase (TPI), enolase (ENO), phosphogluconate dehydrogenase (GND), and aconitase A (ACONTa) in the central carbon metabolism (Figure 3). This is because the relative activity of the oxidative tricarboxylic acid (TCA) cycle, the glyoxylate shunt, and both the oxidative and the non-oxidative pentose phosphate pathway (PPP) change between FDPs. Since PGI and FUM are the only two reactions changing directionalities amongst the four FDPs, it is reasonable to expect the most affected fluxes of reactions to be in their topological vicinity, which is true for GND, TPI, ACONTa, and succinate dehydrogenase (SUCDi) (Figure 3). However, we found large changes in flux magnitudes across the FDPs that were associated with reactions farther away from FUM and PGI, such as the electron transport chain (ETC) reactions, NADH dehydrogenase (NADH16pp) and NAD transhydrogenase (NADTRHD). The TVA studies explain this as the ETC compensates in FDPs 2-4 for producing NADPH (Supp 3 and Supp 4). Additionally, the RSS flux value for GND was considerably smaller in FDP1 than in the other FDPs, resulting in reduced NADPH production via the oxidative branch of the PPP that is coupled with the ETC (Supp 3). For further comparative TVA studies of the FDPs, we refer the reader to supporting information (Supp 4).

**Figure 3.**
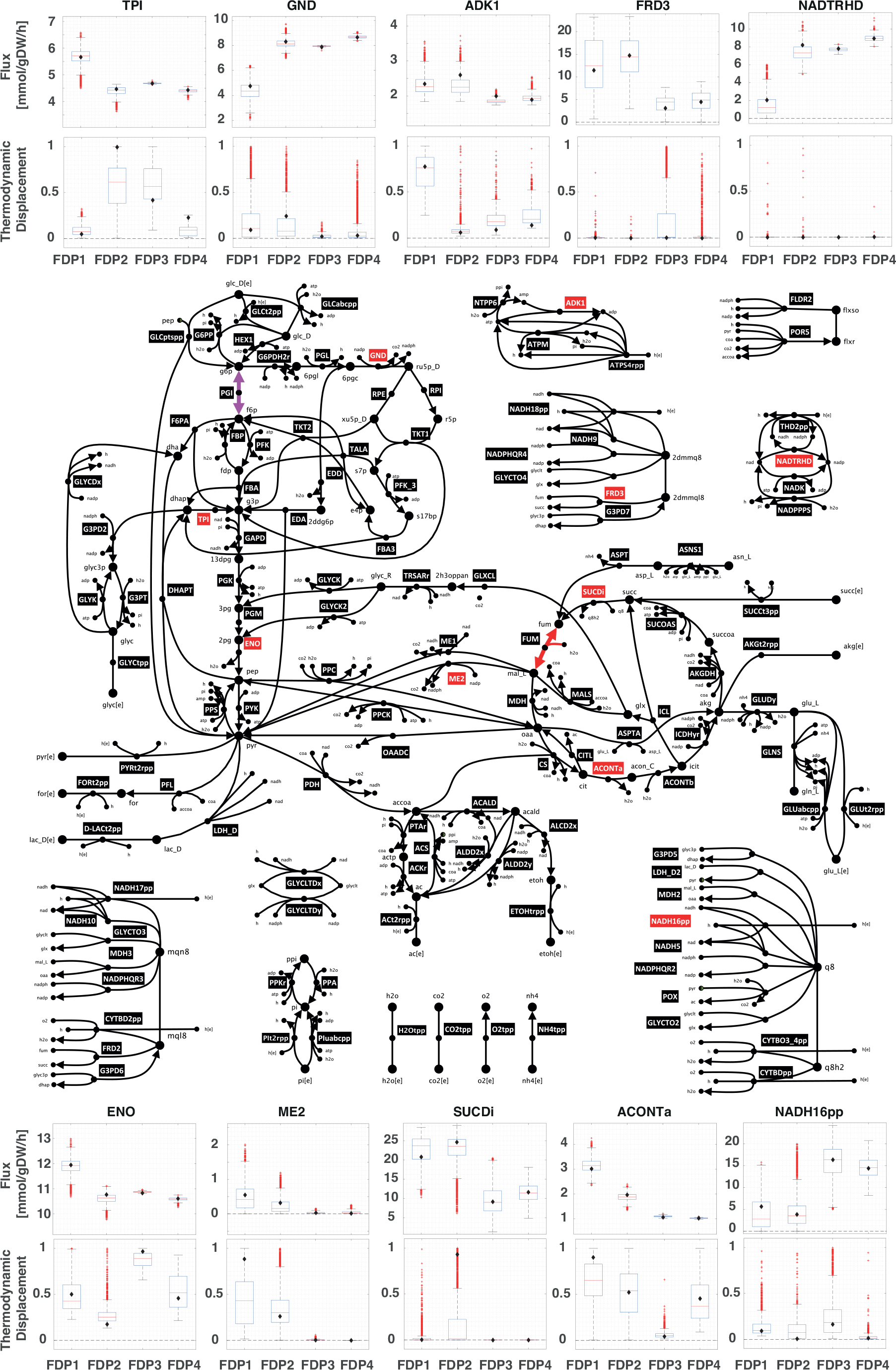
Optimally grown aerobic *E. coli* metabolic network. Each of the 10 reactions labeled in red has an associated graph with the respective flux and thermodynamic displacement distributions for each FDP. The edges of the red boxes represent the thermodynamically feasible bounds for flux and thermodynamic displacement. The boxplots show distributions for 5,000 samples. The central red line indicates the median, and the bottom and top edges of the box indicate the 25^th^ and 75^th^ percentiles, respectively. The whiskers correspond to approximately ± 2.7 σ, the standard deviation, or 99.3% coverage if the data are normally distributed. Outliers are the points not covered by the range of the whiskers and are plotted individually using the ‘+’ symbol. The black diamond is the RSS value. Full enzyme and metabolite names are given in supplementary materials (Supp 2).

The differences in the RSS concentration vectors across the FDPs translate into distinctive distributions of the Gibbs free energy across the networks for each FDP. The metabolite concentration values in the RSSs varied the most across the FDPs for the reaction cofactors NAD^+^, NADH, NADP, AMP, and ATP (Supp 3). We also noticed significant differences in some central carbon metabolite RSS concentrations, such as: 6-phospho-D-gluconate, D-glucose-6-phosphate, D-fructose-6-phophate, D-xylulose 5-phosphate, sedoheptulose 7-phosphate, D-erythrose 4-phosphate, phosphoenolpyruvate, fumarate, L-malate, citrate, and oxaloacetate (Supp 3). These metabolites and the aforementioned cofactors participate in most of the network reactions, causing the thermodynamic displacements of reactions including GND, NADH16pp, SUCDi, adenylate kinase (ADK1), and ME2 to change considerably across RSSs of the FDPs (Figure 3). As observed for the RSS fluxes, reactions that were either topologically close to the bidirectional reactions FUM and PGI and some topologically distant reactions in the ETC displayed the most considerable changes in thermodynamic displacement (Figure 2-3). It is particularly important to recognize that the change in directionality of one reaction between two FDPs can actually cause thermodynamic displacements to change across the whole metabolic network (Figure 4). Hence, the FDP affects the Gibbs free energy distribution across the network, which in turn can affect a greater part of the network than just the topological neighbors of the bi-directional reactions that change directionality between FDPs.

**Figure 4.**
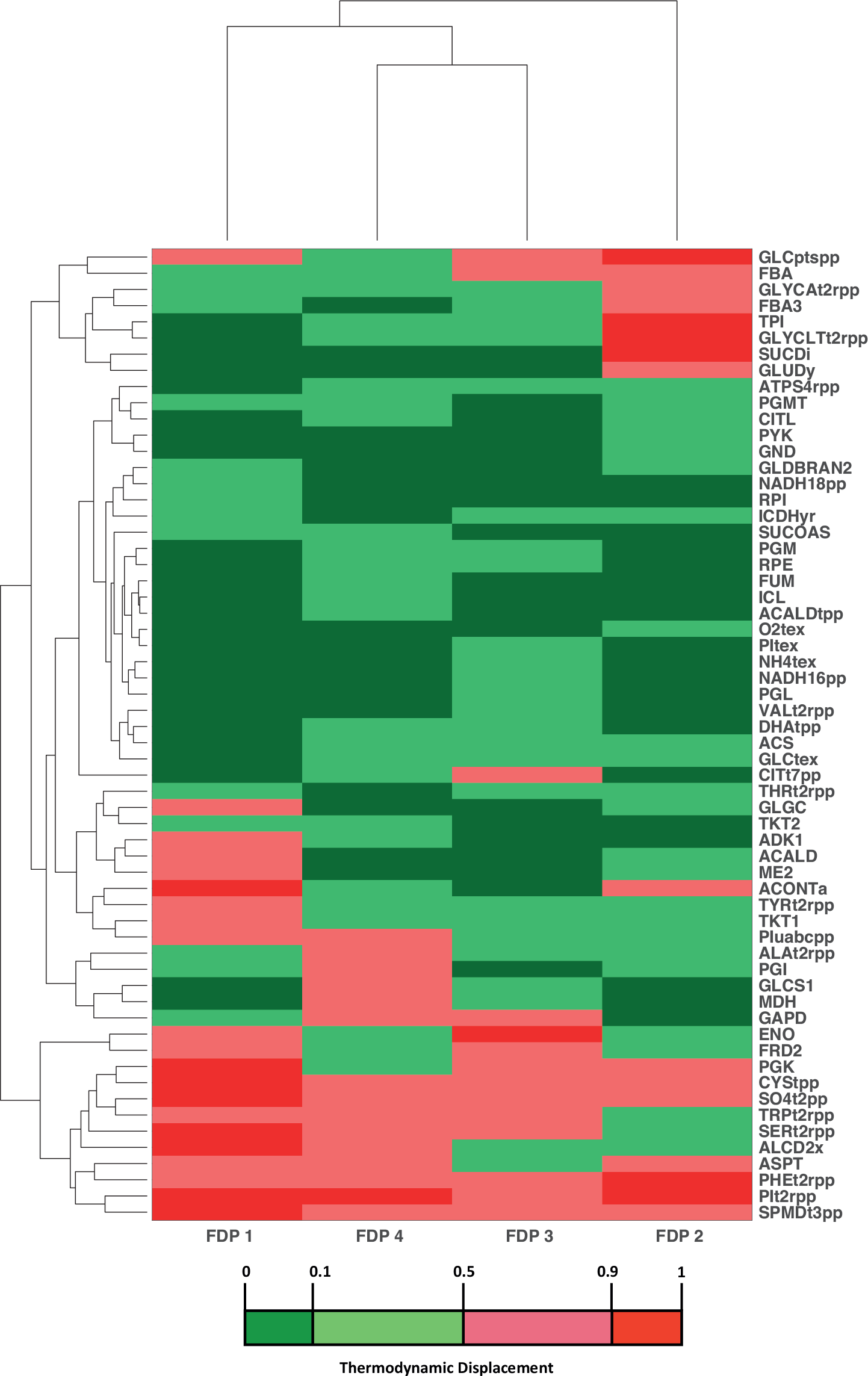
Cluster analysis of thermodynamic displacements across FDPs. Heat map showing thermodynamic displacements of the reactions across each FDP. Reactions differed the most in thermodynamic displacement across FDPs based on categorization of displacements (Methods). The rows represent the similarity between reactions and the columns represent the similarity between FDPs. The distances between the dendrograms were computed based on the Euclidean distance between the thermodynamic displacements, both column- and row-wise. Full enzyme names are given in supplementary materials (Supp 2).

#### 2.2.2. Analysis of control patterns

Controlling the levels of various enzymes in a target organism can help to achieve the desired levels of bioengineered products or metabolites. Determining the degree of control of various enzymes in each FDP can help find the key locations to target, and we did this by sampling the kinetic parameter space uniformly (Methods) using the ORACLE [15–17, 23, 28–30] workflow, generating a population of 50,000 stable kinetic models for each FDP. The kinetic parameter space was sampled based on the degree of saturation of the enzyme active sites, as proposed previously by Hatzimanikatis and colleagues [17]. ORACLE verifies the local stability of the model around the steady state by testing that the Jacobian matrix has no positive real eigenvalues for the sampled set of parameters. We then calculated, for the stable models, the flux control coefficients (FCCs), representing the fold change in a specific flux with respect to the perturbation of an enzyme’s activity, of 275 enzymatic reactions with respect to their enzymes. We then compared the differences in FCCs across FDPs for the populations of stable kinetic models.

If the signs of a FCC are not the same across FDPs, the FCC depends on the FDP, and making metabolic engineering decisions is ambiguous. This means that the alternative steady states have a significant impact on the FCC, and we should be careful when deriving conclusions. FCC values with an absolute mean value larger than 0.1 across all the FDPs have significant control over the fluxes in the network (Methods). Fluxes smaller than 0.01 mmol/gDW/h were not considered, as we focused our analysis around central carbon metabolism. To investigate the differences in control patterns for each FDP, we compared the sign of these FCCs across the FDPs (Figure 5) because the sign determines the increase or decrease in magnitude of a flux upon perturbation of an enzyme level. Hence, the sign can indicate if it may be possible to overexpress, down-regulate, or even suppress a gene to achieve a target enzyme level for bioengineering purposes. If the signs of the mean FCCs are equal across all FDPs, we have consensus, and the FCCs are independent of the FDP. This indicates that the predictive conclusions drawn should be valid for all the tested alternative steady states, suggesting that our metabolic engineering conclusions are more robust. Nearly 75% of the FCCs studied agreed in sign across all the FDPs, meaning that most metabolic flux response predictions are consistent (Figure 5), though the 25% of potentially inconsistent predictions highlights the importance of considering alternative steady states. As we sampled the kinetic parameter space uniformly for the FDPs, differences in the thermodynamic displacement (Figure 4) between these FDPs are the main reason behind these variations in their control pattern. Further discussions around these differences are in the supplementary document (Supp 4).

**Figure 5.**
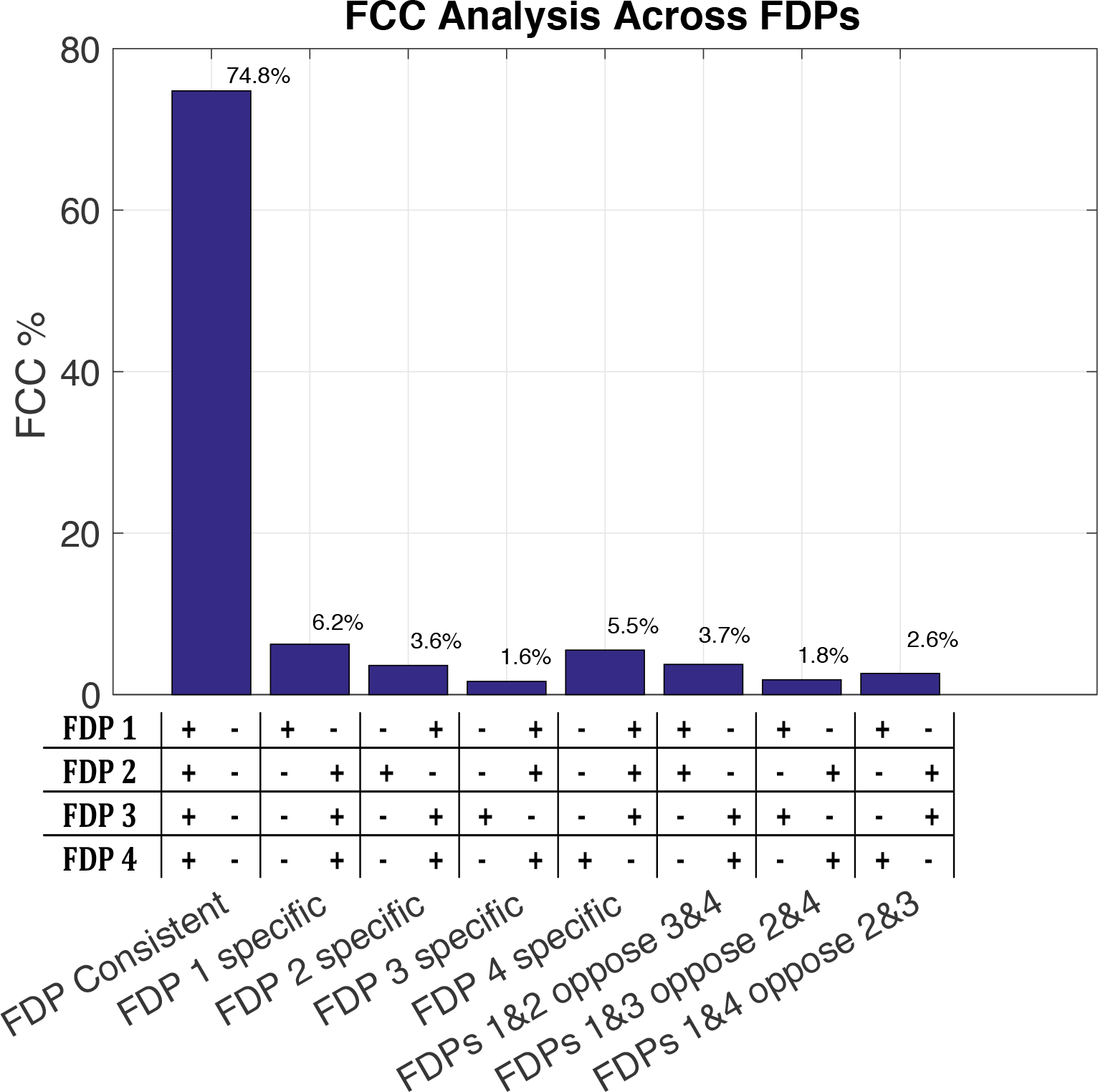
General statistics on FCCs across FDPs. Histogram displaying the fraction of reactions that have FCCs with a certain sign pattern across the FDPs. There are three main categories of the FCCs: (i) consistent among all FDPs, (ii) FDP specific, and (iii) two FDPs contradicting two other FDPs. The FCCs were averaged over the 50,000 samples for each FDP, and the ones selected for analysis had a mean value larger than 0.1 (10% fold change). For example, to assist reading the figure, the column “FDP 1 specific” has two possible scenarios as it contains FCCs that are positive in FDP1 and negative in the other three FDPs as well as FCCs that are negative in FDP1 and positive in the three other FDPs.

#### 2.2.3. Ranking target enzymes for flux control

Some of the fundamental biological tasks performed by a cell include substrate uptake, product excretion, and cellular growth, represented by *μ*. Since we modeled the physiology of optimally grown *E. coli*, we first studied what enzymes have control over cellular growth. These enzymes were considered as attractive target candidates for genetic manipulation to improve cellular growth. We selected the top five enzymes with high absolute FCC values for each FDP and computed the control exerted by these enzymes over cellular growth (Figure 6). Several enzymes such as PGM, RPE, TPI, PPC, and NAD kinase (NADK) had considerably different control patterns for *μ* across FDPs in terms of magnitude and sign. Because of the abovementioned differences in the thermodynamic displacement of enzymes across the FDPs, it was not surprising to see opposing FCC signs. For instance, TPI is far away from equilibrium for FDP1 and near equilibrium for FDP2 (Figure 4), resulting in different conclusions when considering control coefficients of cellular growth (Figure 6). In contrast, PGM is always far away from equilibrium but, due to kinetic coupling, has considerably different levels of control over *μ* across FDPs, indicating the importance of considering alternative steady states. More importantly, we also found enzymes that agreed in terms of sign across the FDPs. NADTRHD, phosphofructokinase (PFK), and ATP maintenance (ATPM) were the top target enzymes - independent of the FDP - for improving the cellular growth of optimally grown *E. coli*.

Because in the studied physiology, growth is based solely on glucose, we decided to study how consistent the FCCs of glucose uptake via D-glucose transport (GLCptspp) were across FDPs. PGM, PFK, RPI, and RPE agreed across all the FDPs in terms of sign and the magnitude of their FCCs, making them attractive metabolic engineering targets for increasing GLCptspp flux (Supp 4, Figure S4). Based on a consistent magnitude across all the FDPs, PGM and PFK were the top two target enzymes that seemed to control the glucose uptake of optimally grown *E. coli*. TPI, PPC, NADTRHD, glucose 6-phosphate dehydrogenase (G6PDH2r), and 6-phosphogluconolactonase (PGL) control GLCptspp in at least one FDP but not across all. As for the control of cellular growth, the differences in thermodynamic equilibrium and kinetic coupling between the FDPs explain these results.

These observations emphasize the importance of considering enzyme kinetics and the existence of alternative steady states before making metabolic engineering conclusions based on kinetic models, especially considering that further similar observations were made for FCCs of other fluxes. Generally, we noticed that the enzymes whose control remained unchanged across FDPs were found in the central carbon metabolism. Inconsistent enzyme control was observed in peripheral and transport reaction enzymes topologically further away from the central carbon pathways.

**Figure 6.**
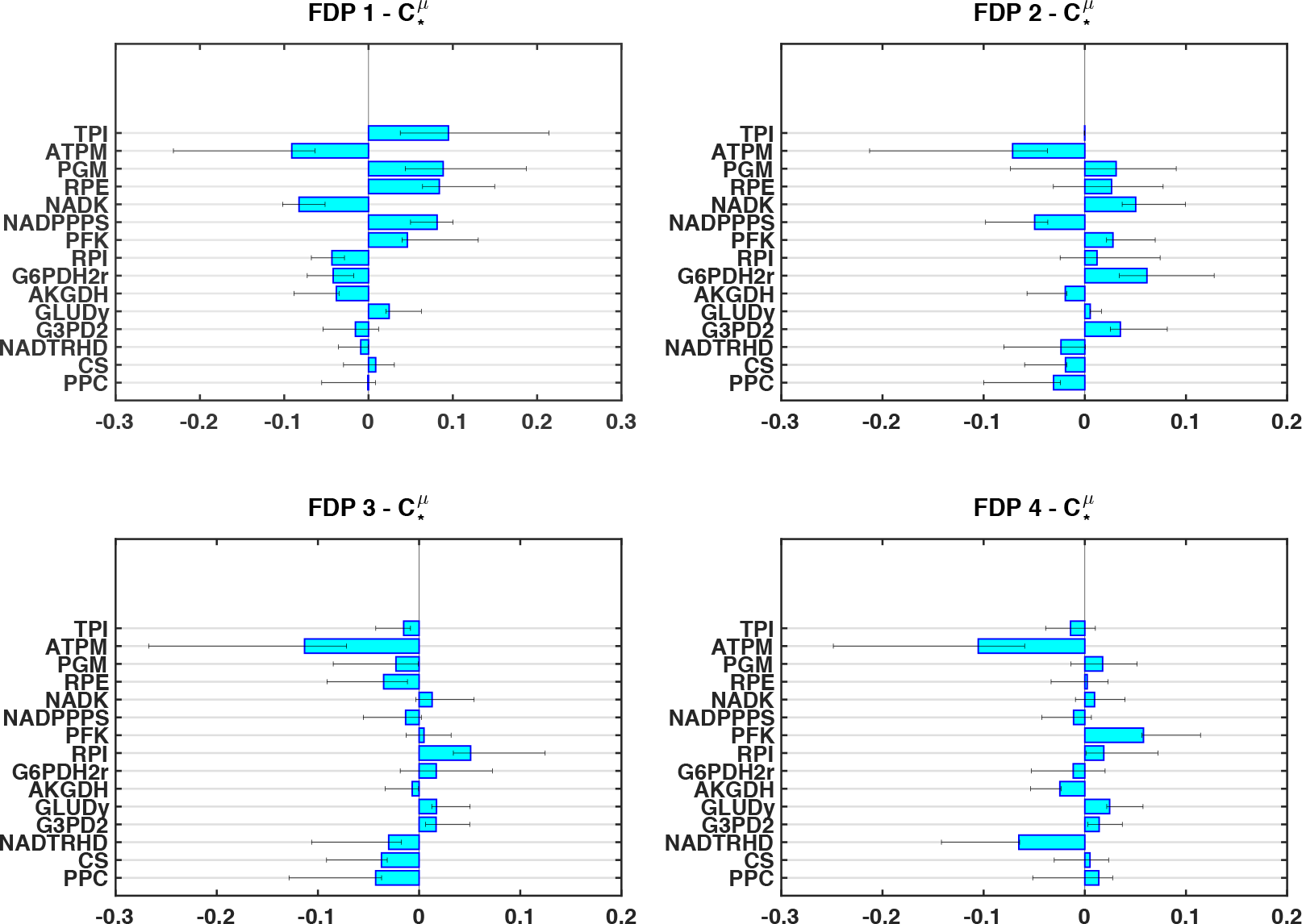
Flux control across FDPs for cellular growth, *μ*. Illustration of the union of the top five enzymes across the FDPs in terms of absolute control over cellular growth. The whiskers correspond to the upper and lower quartiles of the 50,000 FCC populations, and the bars correspond to the means. Full enzyme names are given in supplementary materials (Supp 2).

#### 2.2.4. Study of uncertainty in FCCs

To characterize the variability of cellular growth FCCs with respect to all central carbon enzymes (i.e., no transporter nor exporter) for the populations of 50,000 models (Figure 7), we studied the uncertainty across the four FDPs using PCA (Methods). The first two principal components (PCs) covered a majority of the variance, with 93%, 62%, 56%, and 69% for FDP1-4, respectively. For FDP2-4, at least seven, eight and six PCs, respectively, were required to account for more than 90% of the variance between the FCC populations. This suggests that the uncertainty in the cellular growth FCCs was considerably more distributed for FDP2-4 than for FDP1 that required only two PCs.

In PCA, each variable has a score on the PCs that are under consideration, which corresponds to its contribution to the variability described by the given PC. In FDP1, the growth FCCs with respect to the enzymes NADK and NADPPPS corresponded to the highest PC scores in terms of magnitude along the first PC, suggesting that most of the uncertainty comes from these ETC enzymes (Figure 7). Their scores on both PCs were strongly opposed in terms of sign, suggesting that these FCCs anti-correlate. In fact, the cellular growth FCCs of NADK and NADP phosphatase (NADPPPS) had a −1.00 Pearson correlation coefficient, further indicating that they were exactly anti-correlated. On the other hand, enzymes PGM and RPM had very similar PC scores, and we note that the correlation coefficient of these FCCs was 0.91, indicating a near-perfect correlation.

Similarly to FDP1, we studied FDP2-4 to find underlying covariance patterns between the cellular growth FCCs (Figure 7). We noticed that certain trends were preserved between the FDPs as, for instance, the PCA scores of NADPPPS and G6PDH2r tended to have an opposing sign across the four FDPs for at least one of the plotted PCs. In fact, NADPPPS and G6PDH2r cellular growth FCCs had Pearson correlation coefficients of −0.81, −0.84, −0.53, and −0.71 for FDP1-4, respectively. Hence, we can use PCA to explore and unravel covariance patterns in FCCs to understand their underlying functional relationships. Although fully describing the relationship between FCCs remains a non-trivial task, PCA makes strides towards interpreting the various sources of uncertainty.

NADTRHD, PFK, and ATPM were the top candidates for improving cellular growth, as determined previously based on absolute means (Figure 6). If we had to select one of these three enzymes for genetic engineering, we want it to be the one with the least uncertainty. We observed that PFK scores lower than NADTRHD and ATPM on the PCs across all the FDPs (Figure 7), demonstrating the least uncertainty and suggesting that it could be the most prominent target enzyme. A similar analysis could be performed for FCCs related to other reactions, such as glucose uptake (Supp 4, Figure S5). We conclude that to improve growth of aerobically grown *E. coli*, PFK would also be a top candidate enzyme to metabolically engineer, despite the uncertainties.

The uncertainty in the kinetic parameters and its impact on our studies remains difficult to quantify due to the underdetermined nature of this highly non-linear solution space but it could be further characterized by methods such as the one developed by Andreozzi *et al* [22]. For our study, when we next compare the effect of uncertainties stemming from the flux and the concentration steady-state solutions, we decided to fix the distributions of the sampled enzyme saturations to ones obtained from a previous RSS using beta distributions (Methods). This allowed us to keep the same level of uncertainty in all the kinetic parameters for our comparisons.

**Figure 7.**
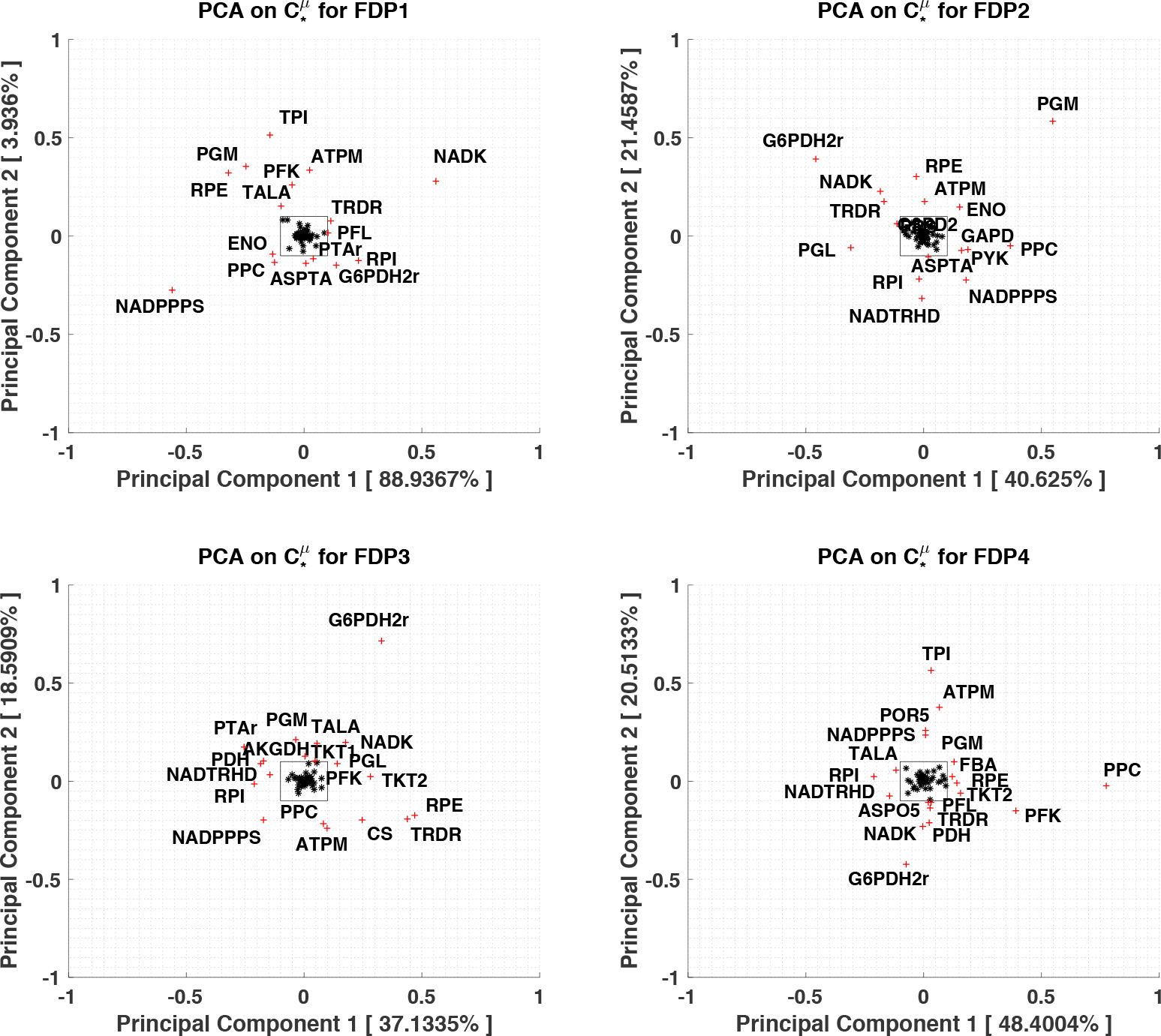
Principal component analysis (PCA) of FCCs of cellular growth across FDPs. PCA was carried out on the cellular growth FCCs of the 50,000 samples for each FDP. Only cellular growth FCCs with respect to non-transporter and non-exporter reactions were considered. The two principal components, namely PC1 and PC2, were plotted to study the variance in the FCC samples. The values in brackets correspond to the variance covered by the principal components (PCs). Full enzyme names are given in supplementary materials (Supp 2).

### 2.3. Impact of flux and concentration profiles

We next assume that we know the directionality of each reaction in our network but the system still remains underdetermined and we have multiple feasible flux and concentration steady states within the allowable solution space. Because of this, we then studied how the underlying uncertainty that results in alternative steady states within a FDP affected the predictions of kinetic models. For this analysis, we selected FDP1 because it has: (1) reaction directionalities corresponding to the more frequently observed *E. coli* wildtype operational state of glycolysis and TCA cycle [36–39], (2) the largest flux variability score, and (3) the highest specificity in control (Figure 5). FDP1 was then more exhaustively sampled with 100,000 iterations of concentrations and fluxes. We chose RSSs from the flux and concentration samples as previously done with the four FDPs, and we used PCA to determine four extreme steady states (ESSs) for the concentrations and four ESSs for the flux solution spaces (Methods). The ESSs are samples with the most distant behavior from the “average” displayed by the RSS. An ESS is a steady state that is located along a PC at one of its extremes and can be used to characterize the “extreme” behaviors of the FDP1.

To study the impact of flux and concentration ESSs on MCA outputs, we had to decouple their effects (Methods), so we isolated the effects of flux and concentration separately in our analysis. Therefore, when we studied the effect of flux, we kept the same concentration RSS and paired it with the four flux ESSs, meaning that we had four pairs of flux ESSs with the same RSS concentration. Similarly, when we decoupled the effect of concentration, we paired the flux RSS with the four concentration ESSs. Therefore, we had a total of eight extreme pairs of flux and concentration steady states to study. We compared these extreme pairs to the reference case, where we had the flux and concentration RSSs paired. For the reference case, we sampled the saturation state space for 50,000 stable kinetic models. We used the distributions of the kinetic parameters from this reference case to generate models for the ESSs (Methods). We sampled 50,000 stable kinetic models for each ESS and computed the FCCs using MCA. We performed a comparative analysis like the comparative analysis of the FDPs to assess the degree of confidence of our conclusions with respect to both the extreme flux profiles and the extreme concentration profiles.

#### 2.3.1. Flux uncertainty propagation to control

Like the comparison of FDPs, the ESS flux profile magnitudes mainly differed in peripheral fluxes, such as glutamate transport, glycogen metabolism, and ETC reactions (Supp 3). Noticeable differences in the central carbon fluxes greater than 1 mmol/gDW/h were seen in pyruvate kinase (PYK), fumarate reductase (FRD3), ME2, NADH17pp, NADH18pp, and NADTRHD. To assess how this variability in fluxes affected the degree of confidence in our MCA conclusions, we considered control over glucose uptake and cellular growth. For glucose uptake, the top enzymes of the flux ESSs, aspartate transaminase (ASPTA), PFK, AKGDH, TKT1, ENO, and TPI, are reasonable candidates for improving uptake because they are all qualitatively in agreement across the flux ESSs and have a control value larger than absolute 0.1, indicating significant control over their networks. This excludes PDH (Figure 8A). Citrate synthase (CS) and G6PDH2r, may appear to be attractive targets based on some of the ESSs, but since this property is not in consensus agreement across all ESSs, they are less reliable targets. The top enzymes controlling cellular growth were sensitive to the ESSs, as just 47% of them were in agreement sign-wise with each other (Supp 4, Figure S6). However, we can still find reliable target enzymes, such as ENO, AKGDH, and glutamate dehydrogenase (GLUDy), mainly in the central carbon reactions that have a reasonable magnitude and consensus agreement.

**Figure 8.**
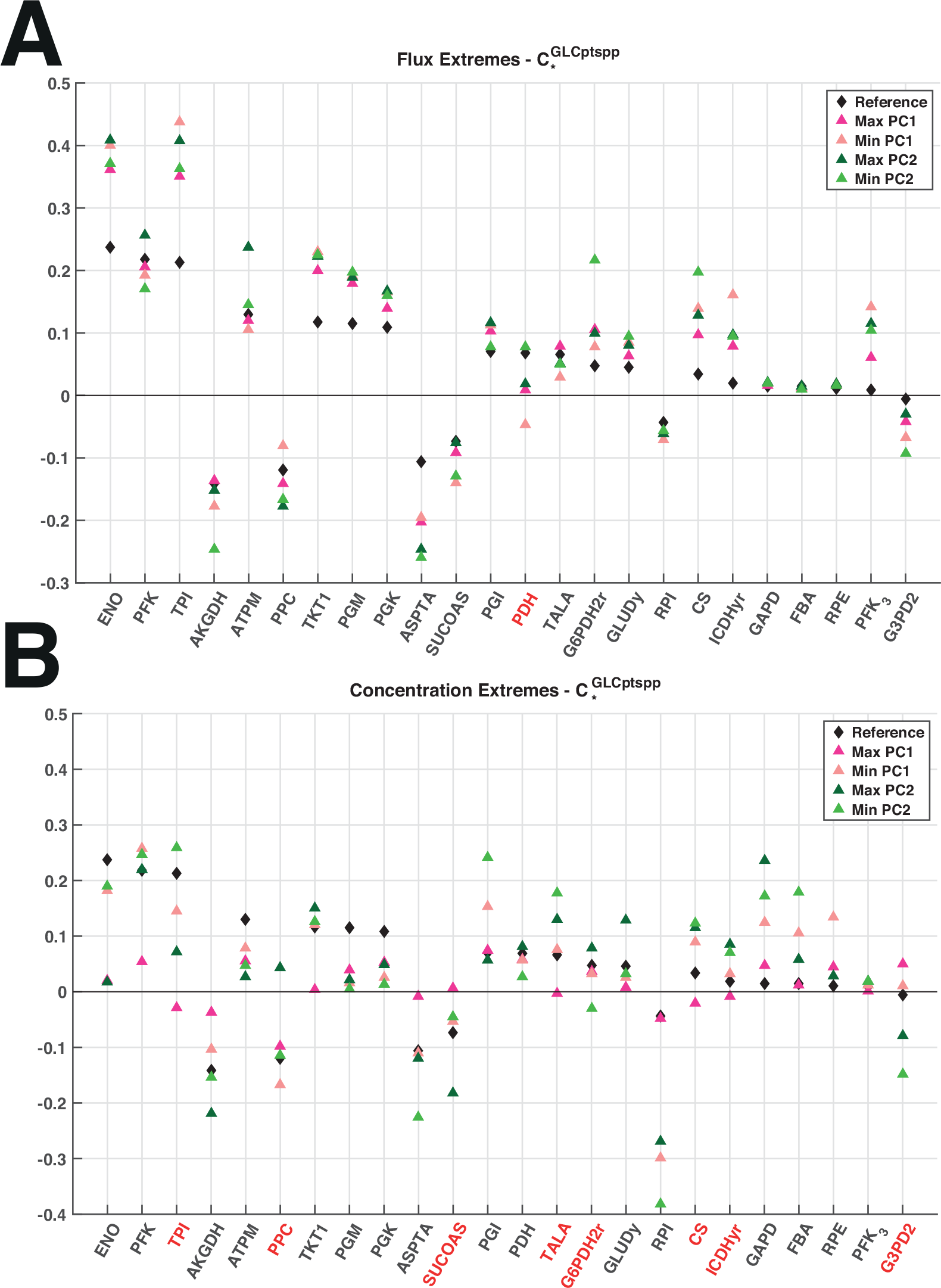
Flux control patterns across extreme steady-state solutions. Illustration of the union between the top 10 enzymes across the **(A)** flux and the **(B)** concentration ESSs in terms of absolute control over glucose uptake. FCCs were sorted in decreasing order of absolute magnitude of the RSSs (reference). The enzyme names in black indicate that the FCCs were sign-consistent (in agreement) and red if they were sign-inconsistent (in opposition). Full enzyme names are given in supplementary materials (Supp 2).

#### 2.3.2. Concentration uncertainty propagation to control

We repeated the previous analysis for ESS concentration vectors to study how they impact MCA outputs. The main concentration differences between the extreme concentration vectors were for amino acids (R-glycerate, L-glutamine, L-lysine, D-alanine, L-proline), inorganics (potassium, iron, and cobalt), cofactors (NAD and AMP), and several biomass building blocks (Supp 3). We considered the top enzymes for glucose uptake FCCs and noticed that the MCA conclusions were more sensitive to concentration values (Figure 8B) than to variations in flux values (Figure 8A). Candidate target enzymes to improve glucose uptake would be PFK, GAPD, ENO, and RPI, as at least three out of five of the steady states are consistent and have a control value larger than 0.1 (Figure 8B). Hence, the metabolic engineering decisions derived from the MCA outputs appear to be more sensitive to concentration values rather than flux values. As the biomass building block and amino acid metabolite concentrations were changing between these ESS concentration vectors, it makes sense that they would have a higher FCC variability. The concentration values in turn directly impact the thermodynamic displacements and the enzyme saturation states, which impact the MCA conclusions. The cellular growth FCCs were very sensitive to the ESS metabolite concentration vectors because most them were in sign disagreement (Supp 4, Figure S6). NADTRHD and ATPM were the most appealing enzymes for controlling cellular growth due to sign and magnitude consistency across the ESSs of their FCCs.

## 3. Conclusions

This work studied the impact of alternative concentration and flux steady states on the conclusions derived from the MCA outputs of the non-linear kinetic models built around them using the physiology of optimally grown *E. coli*. We show that different FDPs can lead to distinct metabolic engineering conclusions when analyzing output FCCs of the non-linear models. The ME2, PPC, and PGI examples illustrate how thermodynamics and kinetic coupling can change the control from one FDP to another. These enzymatic reactions were topologically close to the bidirectional reactions that changed between the FDPs, though, less intuitively, we also noticed that there were changes in thermodynamic displacements across FDPs in enzymes that were topologically far away from the bidirectional reactions. We then studied the uncertainty within a single FDP, and using PCA to study the extremes of the solution space, found that within a FDP, MCA outputs appeared to be more sensitive to concentration values rather than flux values. These observations emphasized the importance of considering alternative solutions when studying a physiology as the steady state affects directly the decisions for hypothesis generation in basic research and design in synthetic biology and metabolic engineering. Hence, we propose a workflow for assessing this uncertainty to make more reliable metabolic engineering decisions that can be broadly applied to any kinetic model to improve the predictions resulting from it.

We then used our workflow to pick target enzymes for genetic modification, identifying NADTRHD, PFK, and ATPM as the top target enzymes independent of the FDP for improving the cellular growth of optimally grown E. coli. PFK and PGM were selected as top enzymes independent of the FDP for improving glucose uptake of optimally grown E. coli. We stress the importance of selecting target enzymes that exhibit control across all the FDPs to make more reliable decisions, highlighting the need to consider alternative steady states when building non-linear kinetic models for a given physiology, as they have imminent implications on the conclusions derived from the MCA. The herein proposed workflow can be used to suggest metabolic engineering decisions for a given study and can provide insights into the design of experiments, as the ranking of candidate enzymes can highlight reactions or enzymes that need further characterization and study due to their variability.

## 4. Methods

### 4.1. Reduced *E. coli* model

The model stoichiometry for this study was derived from *E. coli* iJO1366 [25] using redGEM, a systematic framework for developing core models that are consistent with their genome-scale counterparts [26, 27]. The resulting reduced models are context-specific and in the process of reduction it is important to define the carbon sources, the content of media and also the metabolic subsystems of interest for the study. We applied the redGEM algorithm on the latest genome scale model. We used a minimal media with glucose as the sole carbon source and the selected starting subsystems were ones pertaining to central carbon metabolism (glycolysis/gluconeogenesis, citric acid cycle, pentose phosphate pathway, pyruvate metabolism, and glyoxylate metabolism). Omics data for the physiology of optimally grown *E. coli* under aerobic conditions were extracted (Supp 1) from McClosekey *et al.*[24]. The data were integrated in the form of constraints into the MILP formulation of the thermodynamics-based metabolic flux analysis [35].

We make the following directionality assumptions for several bi-directional reactions:

1. Fructose-biphosphate aldolase (FBA) that is part of mid-lower glycolysis is set towards catabolism [31].
2. The bi-directional transports of magnesium and phosphate are both set towards uptake [32, 33].
3. Acetate kinase (ACKr) and phospho-transacetylase (PTAr) are both set towards the acetate production, because acetate is one of the main by-products [24].
4. The succinyl-CoA synthetase (SUCOAS) is set towards the production of succinate [24].

The polyphosphate kinases (PPK2r, and PPKr) are set towards the polyphosphate polymerization [34].

For some of intracellular metabolites, a corresponding transport reaction has not been biochemically characterized and does not appear in the *E. coli* iJO1366 and in our reduced model. However, these metabolites, unless they are highly polar or very large, are subject to passive diffusive transport through the cell membrane. Therefore, we explicitly added transport reactions for these metabolites that operate at least at basal level (10^-6 mmol/(gDW*h)).

### 4.2. Identification of alternative flux directionality profiles

As first step (Figure 1), in order to identify the reactions that are able to operate in both directions, flux variability analysis (FVA) was performed [40]. If the system has a number *z* of bi-directional reactions, it could have up to 2^*z*^ FDPs as in a FDP, reactions can only operate in a unique direction. We enumerated the FDPs by adjusting the boundaries of the bi-directional reaction so that they can only operate in a unique direction. We define the coefficient of variability, *CV*_*i*_, as:

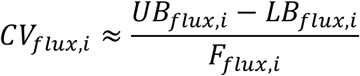

where, *UB* and *LB* are the upper and lower bounds respectively of the flux *i* derived using thermodynamic-based variability analysis (TVA)[7, 35]. *F* is the average of *UB* and *LB*. We define the flux variability score of each FDP as the Euclidean norm of the vector whose entries are the *CV* of each flux. The FDP with the highest flux variability score has the highest relative flexibility in terms of the allowable flux ranges.

### 4.3. Computation of reference and extreme steady states for alternative FDPs

For each of the identified FDPs, in step two (Figure 1), we sample the solution space of concentrations and fluxes without violating physiological, thermodynamic and directionality constraints. The convexity of these solution spaces enables us to efficiently generate sets of flux and concentration samples using the Artificial-Centering Hit-and-Run sampler in the COBRA Toolbox [41–43]. We perform principal component analysis (PCA) on the generated samples to select reference and extreme samples [44]. The first seven principal components (PC) were used as for the fluxes and the concentrations they covered above 90% of the sample variance. The reference sample is chosen so that its vector projections onto the seven PCs are minimal. We get the two extreme samples of a PC, PCmax and PCmin, by respectively finding the 0.1% top and the 0.1% bottom samples based on their magnitude of vector projections onto the given PC. Out of the 0.1% top and the 0.1% bottom samples we chose the samples that have the smallest magnitude of vector projections onto the other PCs (Supp 4).

### 4.4. Analysis of alternative solutions between FDPs

#### 4.4.1. Thermodynamic displacement analysis

Within each FDP, in step 3 (Figure 1), we compute the displacement of the reactions from thermodynamic equilibrium, Γ [17, 45, 46]. For a simple uni-uni reaction with a substrate *S* and a product *P*, the thermodynamic displacement, *Γ*, is defined as:

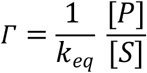

where, *k*_*eq*_ is thermodynamic equilibrium. More specifically, we first use the vector of the reference steady-state concentrations together with values of standard Gibbs free energies of reactions to compute *Γ*. For reactions with negative Gibbs free energy, 0 < *Γ* < 1. For reactions that are far away from equilibrium *Γ* is close to 0, and for reactions near equilibrium *Γ* ≈ 1. We then classify the reactions in terms of *Γ* in the following four classes: reactions that operate (i) near equilibrium (NE), 0.9 ≤ *Γ* ≤ 1; (ii) near to middle equilibrium (NM), 0.5 ≤ *Γ* ≤ 0.9; (iii) middle to far from equilibrium (MF), 0.1 ≤ *Γ* ≤ 0.5; and (iv) far from equilibrium (FE), 0 ≤ *Γ* ≤ 0.1. The information about *Γ* is important, as it is known that enzymes that operate near equilibrium do not have control over fluxes and concentrations in the network [17].

Within the MCA framework, Kaeser and Burns [47] define the concentration control coefficients, 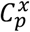, and the flux control coefficients, 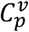, as the fracitonal change of metabolite concentrations and metabolic fluxes, respectively, in response to fractional change of system parameters. According to the log(linear) formalism [48, 49], we can derive 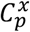 and 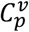 as:

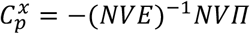

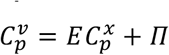

where, *N* is the stoichiometric matrix, *V* is the diagonal matrix whose whole elements are the steady-state fluxes, *E* is the elasticity matrix with respect to metabolites and *Π* is the matrix of elasticities with respect to parameters. If we now consider a uni-uni reaction, *i*, with a substrate *S* and a product *P* we write its reaction rate *v*_*i*_ as follows:

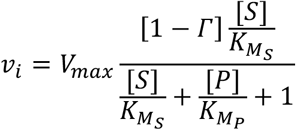

where, *V*_*max*_ is the maximum velocity at enzyme saturation, and, 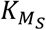 and 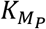, are the Michaelis constants of *S* and *P,* respectively. We define, as done previously by Hatzimanikatis and coworkers [15], the elasticities with respect to *S* and *P*, respectively, as:

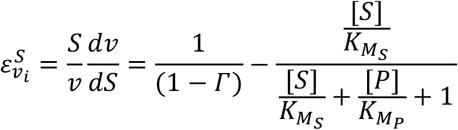

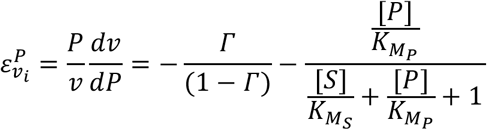

where, 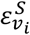 and 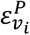 are entries of the elasticity matrix *E*. Evidently, if the reaction is at thermodynamic equilibrium (i.e. *Γ* ≤ 1), the first terms of the elasticity terms 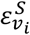 and 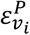 tend towards infinity and we consequently have no control with respect the considered enzyme. However, if the reaction is far away from thermodynamic equilibrium (i.e. *Γ* ≤ 0), the second terms of 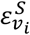 and 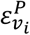 can have impact on the elasticities, potentially resulting in control. Hence, it is essential to consider thermodynamic displacement with the kinetics in order to understand control at systems level. The elasticity matrix *E* is directly affected by *Γ*, and hence the control coefficients 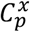 and 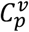 will also be impacted.

#### 4.4.2. Kinetic parameter sampling

We build populations of kinetic models for the computed vectors of the reference steady-state fluxes and concentrations. We integrate the information about the kinetic properties of enzymes available from the literature [50] and the databases [51, 52]. We use the reversible Hill kinetics [53] and convenience kinetics [54] for reactions with unknown kinetic mechanism. For kinetic mechanisms with no or partial information about their parameter values we sample the space of kinetic parameters by direct sampling of the degree of saturation of the active site of an enzyme considering one [17] or multiple enzymatic steps [45]. We then parameterize a population of kinetic models (Supp 5-6), perform consistency verifications [14, 16, 17], and compute the flux and concentration control coefficients [17, 55]. The consistency verifications include a stability test of the model that verifies the Jacobian matrix has no eigenvalues with positive real part for the sampled set of parameters. This test relies on the assumption that the observed RSSs for flux and metabolite concentration are in a stable steady state at the observed time point. For more details about the ORACLE workflow for construction of large-scale kinetic models that are consistent both with thermodynamics and the observed data, the reader is referred to literature [15–17, 22, 23, 28–30, 45].

#### 4.4.3. General statistics on FCCs across FDPs

We computed FCCs of the 275 enzymatic reactions with respect to their 275 enzymes as a quantitative output to compare how our MCA conclusions were consistent across the FDPs. Thus, we calculated the FCCs

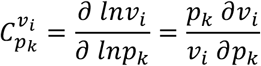

where, *v*_*i*_ is the flux across a reaction *i* and, *p*_*k*_ is the concentration perturbation of an enzyme *k*. We then compute the mean of the FCCs, 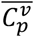, across the kinetic models for an FDP as shown in Table 1.

**Table 1:** Mean of flux control coefficients of a model.

|  |  |  |  |  |
| --- | --- | --- | --- | --- |
| $\overline{C_{p_1}^{v_1}}$ | $\overline{C_{p_2}^{v_1}}$ | $\overline{C_{p_3}^{v_1}}$ | ... | $\overline{C_{p_m}^{v_1}}$ |
| $\overline{C_{p_1}^{v_2}}$ | $\overline{C_{p_2}^{v_2}}$ | $\overline{C_{p_3}^{v_2}}$ | ... | $\overline{C_{p_m}^{v_2}}$ |
| $\overline{C_{p_1}^{v_3}}$ | $\overline{C_{p_2}^{v_3}}$ | $\overline{C_{p_3}^{v_3}}$ | ... | $\overline{C_{p_m}^{v_3}}$ |
| ⋮ | ⋮ | ⋮ | $\ddots$ | ⋮ |
| $\overline{C_{p_1}^{v_n}}$ | $\overline{C_{p_2}^{v_n}}$ | $\overline{C_{p_3}^{v_n}}$ | ... | $\overline{C_{p_m}^{v_n}}$ |

We considered the FCCs for fluxes that were larger than 0.01 mmol/gDW/h across all FDPs because we wanted to focus our study around the reactions that carry more significant amount of carbon (i.e., central carbon metabolism fluxes). Only 126 reactions satisfied this condition, which left us with 34’650 (126 reactions x 275 enzymes) FCCs (Supp 4, Figure S7). To compare more significant FCCs, we only considered ones that had more than absolute 0.1 fold change across the 4 FDPs so that we focus on FCCs with significant control. This meant that we kept 1’263 out of the previous 34’650 FCCs (Supp 4, Figure S8).

### 4.5. Characterizing the distribution of kinetic parameters

#### 4.5.1. Beta distributions

The kinetic parameter solution space is studied in step 4 (Figure 1) by sampling uniformly the degree of saturation of an enzyme’s active site as defined by Wang et al [17]. We obtain distributions of scaled metabolite concentrations from this sampling and consequently, kinetic parameter distributions. The degree of saturation of an enzyme’s active site has a well-defined range from zero to one, allowing us to resort to parametric distributions for their characterization. Beta distributions provide an efficient way of quantitatively expressing variability over a fixed range by estimating its two parameters [56]. These parameters can be obtained and compared for populations of kinetic parameters generated with different operational configurations.

#### 4.5.2. Implying prior beta distributions for sampling

In this work we compare how alternative steady states describing a physiology impact metabolic engineering conclusions. It is thus desirable to ensure that the sampled degrees of saturation of enzyme active sites are similar for the populations of kinetic models built around alternative steady states within FDPs in step 5 (Figure 1). Hence, we compute beta parameters describing the distributions of the kinetic parameters of a given RSS. These beta parameters are used to sample degrees of saturation of enzyme active sites for alternative steady states from similar density distributions using the prior samples. The Beta distribution parameters are implied within the ORACLE workflow as input for sampling degrees of saturations of enzymes when parameterizing new kinetic models. Beta distributions hence bias sample densities for the sampling of degrees of saturation states for an enzyme.

### 4.6. Analysis of alternative solutions within FDPs

We investigate in step 5 (Figure 1) how different flux profiles and metabolite concentration vectors, within FDPs, affect the populations of control coefficients. We separately studied the effects of the flux profiles and the metabolite concentration vectors, in order to decouple their effects on control coefficients. We take the reference steady-state concentration vector and we form the pairs with the extreme steady-state flux profiles computed in step 2 of the procedure (Figure 1). We then generate populations of kinetic models as described in step 3. In the generation of missing kinetic information, we use the distributions of kinetic parameters that have been characterized in step 4 for this FDP. This way, we obtain alternative populations of kinetic models that have in common the reference steady-state concentration and the distribution of kinetic parameters. We compare these populations of kinetic models together with the population of kinetic models that was computed in step 3 for the reference steady state of this FDP. This enables the assessment of the effects of alternative flux profiles within the FDP onto the control coefficients.

The effects of alternative values of concentrations on control coefficients are estimated in an analogous way, where we take the reference steady-state flux and we form the pairs with the extreme steady-state concentrations and we repeat the procedure discussed above. Taken together these two comparisons of alternative solutions allow us to identify sets of enzymes within a FDP whose control over the fluxes and concentrations in the network is robust both with respect to the alternative concentrations and fluxes. We also identify enzymes that are robust only with respect to the alternative concentrations or alternative fluxes.

### 4.7. Metabolic Engineering and Synthetic Biology Design

We next analyze in step 6 the results obtained in steps 3-5 (Figure 1) in the light of metabolic engineering and synthetic biology design. We single out the enzymes whose control over fluxes and concentrations of interest is consistent over all FDPs and within FDPs. In this step we can also design the experiments that would give sufficient information for discriminating alternative solutions between FDPs and within FDPs.

## Acknowledgements

We would like to thank Joana Pinto Vieira for her help with editing this manuscript.

## Supplementary

**S1 Table. Fluxomics and metabolomics data incorporated in the model.** Table giving the fluxomics data in mmol/gDW/h and the concentration data in log(M).

**S2 Table. Thermodynamics-based metabolic flux analysis model.** Spreadsheet giving the list of metabolites, reactions, variables, constraints and compartmentalization in the flux directionality profile 1.

**S3 Table. Thermodynamic variability analysis.** Spreadsheet giving the flux ranges and Gibb’s free energy ranges for reactions and the metabolite log(M) concentration ranges.

**S4 Supplemental Material.** Additional information and supplemental figures.

**S5 Text. System of ordinary differential equations describing FDP1.** Non-linear kinetic model of *E.coli* for FDP1 giving the system of ordinary differential equations.

**S6 Table. Reaction mechanisms describing the systems.**

## References

1. Nielsen J. Systems Biology of Metabolism. Annual Review of Biochemistry. 2017;86(1):245–75. doi:10.1146/annurev-biochem-061516-044757.

2. Alper H, Stephanopoulos G. Engineering for biofuels: exploiting innate microbial capacity or importing biosynthetic potential? Nature Reviews Microbiology. 2009;7:715. doi:10.1038/nrmicro2186.

3. Li G, Wang J-b, Reetz MT. Biocatalysts for the pharmaceutical industry created by structure-guided directed evolution of stereoselective enzymes. Bioorganic & medicinal chemistry. 2017.

4. Blazeck J, Alper H. Systems metabolic engineering: Genome-scale models and beyond. Biotechnol J. 2010;5(7):647–59.

5. Thiele I, Palsson BØ. A protocol for generating a high-quality genome-scale metabolic reconstruction. Nature protocols. 2010;5(1):93–121.

6. Zamboni N, Fendt S-M, Ruhl M, Sauer U. 13C-based metabolic flux analysis. Nat Protocols. 2009;4(6):878–92. doi:http://www.nature.com/nprot/journal/v4/n6/suppinfo/nprot.2009.58_S1.html.

7. Ataman M, Hatzimanikatis V. Heading in the right direction: thermodynamics-based network analysis and pathway engineering. Current Opinion in Biotechnology. 2015;36:176–82.

8. Soh KC, Hatzimanikatis V. Constraining the Flux Space Using Thermodynamics and Integration of Metabolomics Data. Metabolic Flux Analysis: Springer; 2014. p. 49–63.

9. Salvy P, Fengos G, Ataman M, Pathier T, Soh KC, Hatzimanikatis V. pyTFA and matTFA: A Python package and a Matlab toolbox for Thermodynamics-based Flux Analysis. Bioinformatics. 2018;1:3.

10. Ebrahim A, Brunk E, Tan J, O’brien EJ, Kim D, Szubin R, et al. Multi-omic data integration enables discovery of hidden biological regularities. Nature Communications. 2016;7.

11. Lerman JA, Hyduke DR, Latif H, Portnoy VA, Lewis NE, Orth JD, et al. In silico method for modelling metabolism and gene product expression at genome scale. Nature communications. 2012;3:929.

12. Sánchez BJ, Zhang C, Nilsson A, Lahtvee PJ, Kerkhoven EJ, Nielsen J. Improving the phenotype predictions of a yeast genome-scale metabolic model by incorporating enzymatic constraints. Molecular systems biology. 2017;13(8):935.

13. Chen P-W, Theisen MK, Liao JC. Metabolic systems modeling for cell factories improvement. Current Opinion in Biotechnology. 2017;46:114–9.

14. Miskovic L, Tokic M, Fengos G, Hatzimanikatis V. Rites of passage: requirements and standards for building kinetic models of metabolic phenotypes. Current Opinion in Biotechnology. 2015;36:1–8.

15. Chakrabarti A, Miskovic L, Soh KC, Hatzimanikatis V. Towards kinetic modeling of genome-scale metabolic networks without sacrificing stoichiometric, thermodynamic and physiological constraints. Biotechnol J. 2013;8(9):1043–57. Epub 2013/07/23. doi:10.1002/biot.201300091. PubMed PMID: 23868566.

16. Miskovic L, Hatzimanikatis V. Production of biofuels and biochemicals: in need of an ORACLE. Trends in biotechnology. 2010;28(8):391–7.

17. Wang L, Birol I, Hatzimanikatis V. Metabolic Control Analysis under Uncertainty: Framework Development and Case Studies. Biophysical Journal. 2004;87:3750–63.

18. Khodayari A, Maranas CD. A genome-scale Escherichia coli kinetic metabolic model k-ecoli457 satisfying flux data for multiple mutant strains. Nature Communications. 2016;7.

19. Tran LM, Rizk ML, Liao JC. Ensemble Modeling of Metabolic Networks. Biophysical Journal. 2008;95(12):5606–17. doi:https://doi.org/10.1529/biophysj.108.135442.

20. Schuetz R, Kuepfer L, Sauer U. Systematic evaluation of objective functions for predicting intracellular fluxes in Escherichia coli. Molecular Systems Biology. 2007;3:119. doi:10.1038/msb4100162. PubMed PMID: PMC1949037.

21. Almquist J, Cvijovic M, Hatzimanikatis V, Nielsen J, Jirstrand M. Kinetic models in industrial biotechnology–improving cell factory performance. Metabolic engineering. 2014;24:38–60.

22. Andreozzi S, Miskovic L, Hatzimanikatis V. iSCHRUNK–In Silico Approach to Characterization and Reduction of Uncertainty in the Kinetic Models of Genome-scale Metabolic Networks. Metabolic engineering. 2016;33:158–68.

23. Andreozzi S, Chakrabarti A, Soh KC, Burgard A, Yang TH, Van Dien S, et al. Identification of metabolic engineering targets for the enhancement of 1,4-butanediol production in recombinant E. coli using large-scale kinetic models. Metabolic engineering. 2016;35:148–59. doi:10.1016/j.ymben.2016.01.009. PubMed PMID: 26855240.

24. McCloskey D, Gangoiti JA, King ZA, Naviaux RK, Barshop BA, Palsson BO, et al. A model-driven quantitative metabolomics analysis of aerobic and anaerobic metabolism in E. coli K-12 MG1655 that is biochemically and thermodynamically consistent. Biotechnology and bioengineering. 2014;111(4):803–15.

25. Orth JD, Conrad TM, Na J, Lerman JA, Nam H, Feist AM, et al. A comprehensive genome-scale reconstruction of Escherichia coli metabolism-2011. Molecular systems biology. 2011;7(1):535.

26. Ataman M, Hatzimanikatis V. lumpGEM: Systematic generation of subnetworks and elementally balanced lumped reactions for the biosynthesis of target metabolites. PLoS computational biology. 2017;13(7):e1005513.

27. Ataman M, Gardiol DFH, Fengos G, Hatzimanikatis V. redGEM: Systematic reduction and analysis of genome-scale metabolic reconstructions for development of consistent core metabolic models. PLoS computational biology. 2017;13(7):e1005444.

28. Soh KS, Miskovic L, Hatzimanikatis V. From network models to network responses: integration of thermodynamic and kinetic properties of yeast genome-scale metabolic networks. FEMS Yeast Research. 2012;12:129–43.

29. Wang LQ, Hatzimanikatis V. Metabolic engineering under uncertainty. I: Framework development. Metabolic engineering. 2006;8(2):133–41. doi:Doi 10.1016/J.Ymben.2005.11.003. PubMed PMID: ISI:000236053000005.

30. Wang LQ, Hatzimanikatis V. Metabolic engineering under uncertainty - II: Analysis of yeast metabolism. Metabolic engineering. 2006;8(2):142–59. doi:Doi 10.1016/J.Yinben.2005.11.002. PubMed PMID: ISI:000236053000006.

31. Cooper R. Metabolism of methylglyoxal in microorganisms. Annual Reviews in Microbiology. 1984;38(1):49–68.

32. Nelson DL, Kennedy EP. Transport of magnesium by a repressible and a nonrepressible system in Escherichia coli. Proceedings of the National Academy of Sciences. 1972;69(5):1091–3.

33. Rosenberg H, Gerdes R, Chegwidden K. Two systems for the uptake of phosphate in Escherichia coli. Journal of bacteriology. 1977;131(2):505–11.

34. Kumble KD, Ahn K, Kornberg A. Phosphohistidyl active sites in polyphosphate kinase of Escherichia coli. Proceedings of the National Academy of Sciences. 1996;93(25):14391–5.

35. Henry CS, Broadbelt LJ, Hatzimanikatis V. Thermodynamics-based metabolic flux analysis. Biophysical journal. 2007;92(5):1792–805.

36. Park JO, Rubin SA, Xu Y-F, Amador-Noguez D, Fan J, Shlomi T, et al. Metabolite concentrations, fluxes and free energies imply efficient enzyme usage. Nature chemical biology. 2016;12(7):482–9.

37. Toya Y, Ishii N, Nakahigashi K, Hirasawa T, Soga T, Tomita M, et al. 13C-metabolic flux analysis for batch culture of Escherichia coli and its pyk and pgi gene knockout mutants based on mass isotopomer distribution of intracellular metabolites. Biotechnology progress. 2010;26(4):975–92.

38. Crown SB, Long CP, Antoniewicz MR. Integrated 13 C-metabolic flux analysis of 14 parallel labeling experiments in Escherichia coli. Metabolic engineering. 2015;28:151–8.

39. Fong SS, Nanchen A, Palsson BO, Sauer U. Latent pathway activation and increased pathway capacity enable Escherichia coli adaptation to loss of key metabolic enzymes. Journal of Biological Chemistry. 2006;281(12):8024–33.

40. Mahadevan R, Schilling C. The effects of alternate optimal solutions in constraint-based genome-scale metabolic models. Metabolic engineering. 2003;5(4):264–76.

41. Schellenberger J, Que R, Fleming RMT, Thiele I, Orth JD, Feist AM, et al. Quantitative prediction of cellular metabolism with constraint-based models: the COBRA Toolbox v2.0. Nat Protocols. 2011;6:1290–307. doi:10.1038/nprot.2011.308.

42. Becker SA, Feist AM, Mo ML, Hannum G, Palsson BO, Herrgard MJ. Quantitative prediction of cellular metabolism with constraint-based models: the COBRA Toolbox. Nat Protocols. 2007;2:727–38. doi:10.1038/nprot.2007.99.

43. Kaufman DE, Smith RL. Direction choice for accelerated convergence in hit-and-run sampling. Operations Research. 1998;46(1):84–95.

44. Jolliffe I. Principal Component Analysis. Wiley StatsRef: Statistics Reference Online: John Wiley & Sons, Ltd; 2014.

45. Miskovic L, Hatzimanikatis V. Modelling of uncertainties in biochemical reactions. Biotechnology and Bioengineering. 2011;108:413–23.

46. Heinrich R, Schuster S. The regulation of cellular systems. New York; London: Chapman & Hall; 1996.

47. Kacser H, Burns J, editors. The control of flux. Symp Soc Exp Biol; 1973.

48. Hatzimanikatis V, Floudas CA, Bailey JE. Analysis and design of metabolic reaction networks via mixed-integer linear optimization. AIChE Journal. 1996;42(5):1277–92.

49. Reder C. Metabolic control theory: a structural approach. Journal of theoretical biology. 1988;135(2):175–201.

50. Segel IH. Enzyme Kinetics. 1975.

51. Schomburg I, Chang A, Placzek S, Sohngen C, Rother M, Lang M, et al. BRENDA in 2013: integrated reactions, kinetic data, enzyme function data, improved disease classification: new options and contents in BRENDA. Nucleic Acids Res. 2013;41(Database issue):D764–72. Epub 2012/12/04. doi:10.1093/nar/gks1049. PubMed PMID: 23203881.

52. Wittig U, Kania R, Golebiewski M, Rey M, Shi L, Jong L, et al. SABIO-RK-database for biochemical reaction kinetics. Nucleic Acids Res. 2012;40(D1):D790–D6. doi:Doi 10.1093/Nar/Gkr1046. PubMed PMID: ISI:000298601300118.

53. Hofmeyr J, Cornish-Bowden A. The reversible Hill equation: how to incorporate cooperative enzymes into metabolic models. Comput Appl Biosci. 1997;13:377–85.

54. Liebermeister W, Klipp E. Bringing metabolic networks to life: convenience rate law and thermodynamic constraints. Theoretical Biology and Medical Modeling. 2006;3(41). doi:doi:10.1186/1742-4682-3-41.

55. Hatzimanikatis V, Bailey JE. MCA has more to say. Journal of Theoretical Biology. 1996;182(3):233–42.

56. Hahn GJ, Shapiro SS. Statistical Models in Engineering: Wiley; 1994.

